# The Virulence Index: A Metric for Quantitative Analysis of Phage Virulence

**DOI:** 10.1101/606350

**Authors:** Zachary Storms, Matthew R. Teel, Kevin Mercurio, Dominic Sauvageau

**Affiliations:** Chemical and Materials Engineering, University of Alberta, Edmonton, AB, Canada

**Author notes:** **Correspondence:** Dominic Sauvageau. 1-780-492-8092.

**Keywords:** bacteriophage infection, bacterial reduction curve, comparative virulence, virulence quantification, quality control, high-throughput analysis

## Abstract

**Background:** One of the main challenges in developing phage therapy and manufacturing phage products is the reliable evaluation of their efficacy, performance and quality. Since phage virulence is intrinsically difficult to fully capture, researchers have turned to rapid but partially inadequate methods for its evaluation.

**Materials and Methods:** The present study demonstrates a standardized, quantitative method to assess phage virulence based on three parameters: the Virulence Index (*V*_*P*_) – quantifying the virulence of a phage against a host, the local virulence (*v*_*i*_) – assessing killing potential at given MOIs, and *MV*_*50*_ – the MOI at which the phage achieves 50% of its maximum theoretical virulence. This was shown through comparative analysis of the virulence of phages T4, T5 and T7.

**Results:** Under the conditions tested, phage T7 displayed the highest virulence, followed by phage T4 and, finally, phage T5. The impact of parameters such as temperature and medium composition on virulence was shown for each phage. The use of the method to evaluate the virulence of combinations of phages – e.g. for cocktail formulation – is also shown with phages T5 and T7.

**Conclusions:** The method presented provides a platform for high-throughput quantitative assessment of phage virulence and quality control of phage products. It can also be applied to phage screening, evaluation of phage strains, phage mutants, infection conditions and/or the susceptibility of host strains, and the formulation of phage cocktails.

## Introduction

Despite the significant impact bacteriophages (phages) have had in understanding genetic and gene regulation ^1^, and the enormous estimated number of phages present on this planet ^2^, a relatively small number of phage species have been identified and fewer have been fully characterized. But this is rapidly changing. Initiatives have been launched to both isolate new phage species and annotate the staggering amount of genomic data collected in the growing number of screening and characterization studies (the PhAnToMe project, the Marine Phage Sequencing Project, and ^3,4^ are just some examples). There are also numerous ongoing efforts to identify phages suitable for phage therapy ^5-8^, biocontrol ^9,10^, prevention of biofilms ^11-12^, detection and diagnostics ^13-15^, and as active or structural elements in biomaterials.^16,17^ With the rapidly growing number of applications comes an increasing need for the manufacture of phages and phage products, and by extension approaches and methods to reliably assess their efficacy and quality. As the efficacy of phage products relies on the phage properties, in many ways assessment of quality is tied to phage characterization.

Several standardized growth-associated parameters are used to characterize phages. These include burst size (virus particles produced per infection), eclipse period (period from infection to the production of the first viable intracellular phage virion), latent period (period from infection to cell lysis), adsorption rate, adsorption efficiency, and overall growth rate or ‘phage fitness,’ to name a few. Together, these parameters all contribute to phage virulence, “the killing ability of the phage”, but there is no standardized way to report, or even a clear definition of, phage virulence itself.

In epidemiology, a common measurement of virulence is the reproduction number (*R*_0_) – defined as the average number of additional hosts that the virus will spread to after infecting a single host ^18^. An equivalent measurement is not applicable to the virulent phage life cycle. The number of phage progeny released per infected cell is given by the burst size; when replicating in a healthy, densely growing bacterial culture, the *R*_0_ of a phage is essentially equal to its burst size. However, burst size alone is not a proper indicator of phage virulence. For example, in a study of the efficacy of phage therapy, virulence was found to be an increasing function of both adsorption rate and burst size.^19^ In fact, even more parameters are needed to fully quantify virulence; these include, amongst others, the latent period, adsorption efficiency, and host cell growth rate.^20^ In this context, virulence can be defined as the ability of a phage to kill or damage a host population.^21^ As pointed out by Hobbs and Abedon ^22^, the concept of virulence is often applied to differentiate phages undergoing lytic rather than lysogenic or chronic infections. But even in the case of comparing strictly lytic phages or various clones of a strictly lytic phage, it is possible to infer different degrees of virulence, despite the inconsistencies in terminology.

Common techniques employed to qualitatively measure virulence are spot tests on agar plates 23 and bacterial reduction curves.^24^ A spot test entails spreading a small sample of a specific phage over a bacterial culture growing in a top-agar lawn. This is a quick way to test susceptibility of a bacterial strain to various phages but does not capture infection dynamics.^25^ A bacterial reduction curve is obtained by infecting a liquid bacterial culture with phages and taking periodic optical density measurements, which will be reduced compared to those of a phage-free control. A typical virulence study consists of generating bacterial reduction curves under various conditions and qualitatively comparing which bacterial curve is ‘reduced’ the most. The advantage of this approach is that it is applicable to any phage cultivable in suspended cultures. The disadvantages are that it is time-consuming, non-standardized and only qualitative in its current incarnation. Examples of bacterial reduction curves are plentiful in the literature.^24,26-34^ While all these experiments rely on the same principle, no standardized or quantitative method has emerged to quantify virulence or facilitate comparisons across conditions and studies.

In one study, four isolated phages specific to *Escherichia coli* O157:H7 were screened against hundreds of *E. coli* strains to gather host range and susceptibility data.^24^ The protocol consisted of performing bacterial infections in 220-μl volumes using 96-well plates. Bacterial cultures at the same optical density were inoculated with phages at initial Multiplicities of Infection (MOIs) ranging from 10^-6^ to 10^2^. Plates were incubated at 37°C for 5 hours and then visually observed for signs of cell lysis. Host cell susceptibility was then categorized based on the minimum MOI required to obtain a culture-wide lysis by visual inspection. This approach offers many improvements over the current mix of non-standardized protocols adopted by different researchers. But it has shortcomings. Firstly, it relies on visual inspection rather than a measureable property. Secondly, it fails to capture the dynamics of infection, instead relying on endpoint measurements, which can be misleading. Thirdly, it defines lytic capability in terms of susceptibility of the host rather than the phage.

Another study introduced the concept of comparing the areas under bacterial infection curves to non-infected bacterial growth curves was introduced as a means of evaluating the conditions at which a phage or a combination of two phages was most efficient at killing its host (infective).^35^ Recently, Xie et al. ^36^ built on this concept to demonstrate how bacterial reduction curves performed in microplates can be used for the high throughput evaluation of host range and phage infectivity. In the study, the authors demonstrated how the comparison of the areas under the curves could be used to semi-quantify the efficacy of a phage against a given host or a range of hosts.

In the present study, we detail a method to measure phage virulence that overcomes the shortcomings faced by traditional methods. We build on the premise that comparisons between infected and non-infected bacterial cultures can be used to quantitatively assess the virulence of a phage or phage mixture – as demonstrated in ^35,36^. This method generates a Virulence Index (*V*_*P*_) for a phage infecting a specific host strain under a given set of environmental conditions. The present study, which compares the virulence of phages T4, T5 and T7 under various conditions, shows the Virulence Index can be used to easily compare and quantify the virulence of diverse phages. This protocol greatly simplifies and improves the reliability of virulence measurements, and provides a platform to quantitatively compare between phages and conditions.

The phage research community and industry require a simple, fast, and standardized way to measure quantitatively phage virulence that takes into account all factors affecting virulence. This will greatly facilitate screening, selection, comparison and quality control of phages and phage products for specific scientific, industrial or therapeutic applications. Such a method can also be used to monitor the impact of mutations or adaptation on virulence, and to establish formulations of phage cocktails for various applications.

## Materials and Methods

### Organisms and Media

Cultures of *Escherichia coli* ATCC 11303 used for experimentation were grown overnight in 10 ml of medium. The media used were Bacto Tryptic Soy Broth (TSB; Becton Dickinson, Sparks, MD) and BBL Brain Heart Infusion (BHI; Becton Dickinson, Sparks, MD). Host cultures were grown in 125-ml Erlenmeyer flasks containing 10 ml of medium, agitated at 150 rpm and incubated at 37°C. Phage species used were phage T4 (ATCC 11303-B4), phage T5 (ATCC 11303-B5), and phage T7 (ATCC 11303-B7). Phage stocks were stored at 4°C in TSB at titers of 4.5×10^8^ pfu/ml, 2.3×10^8^ pfu/ml, and 2.1×10^8^ pfu/ml, respectively.

### Bacterial Reduction Experiments

Overnight cultures (100 μl) were used to inoculate 10 ml of fresh medium in an Erlenmeyer flask, incubated at 37°C and 150 rpm. These cultures were allowed to grow beyond the starting bacterial concentration used for the bacterial reduction experiments (10^8^ cfu/ml). All bacterial reduction curves were generated using 96-well plates with 300-μl well volumes. Phage stocks were serially diluted from a concentration of 10^8^ pfu/ml to 10 pfu/ml in 200-μl volumes. Plates were incubated at 37°C for 30 minutes prior to inoculation to ensure stable temperatures. Cell concentrations were adjusted to 10^8^ cfu/ml for every experiment, unless otherwise indicated, yielding MOIs ranging from 10^-7^ to 1. In this report, MOI refers to the initial MOI at the initiation of phage infection. Considering that the working volume used to generate the reduction curves was 250 μL, MOIs lower than 10^-7^ were not tested since, on average, they would have less than 1 phage per culture. A layout of the microplate used for high throughput evaluation of virulence can be seen in Fig. 1A. Four wells of phage-free bacterial cultures were included on every plate as control experiments, in addition to four media-blanks for reference. Experiments were run with all phage samples growing in parallel in a final volume of 250 μl. Since phages are serially diluted to obtain different MOIs, this set-up can be completed rapidly with a multichannel pipetter. Optical density was measured at 630 nm with a Bio-Tek ELx800 Universal Microplate Reader and the data was recorded using KCJunior software. Optical density measurements were taken immediately after inoculation and then at regular intervals afterwards. Between samples, the plates were covered and placed in an incubator shaker at 150 rpm at the specified experimental temperature (continuous incubation and readings in an incubating plate-reader is also possible, even recommended when available). Measurements were taken until the control cultures reached stationary phase. Once all data was collected, areas underneath the optical density vs. time curves were calculated using the trapezoid rule for each well, from the time of infection to the time corresponding to the onset of stationary phase in the phage-free control.

**Figure 1.**
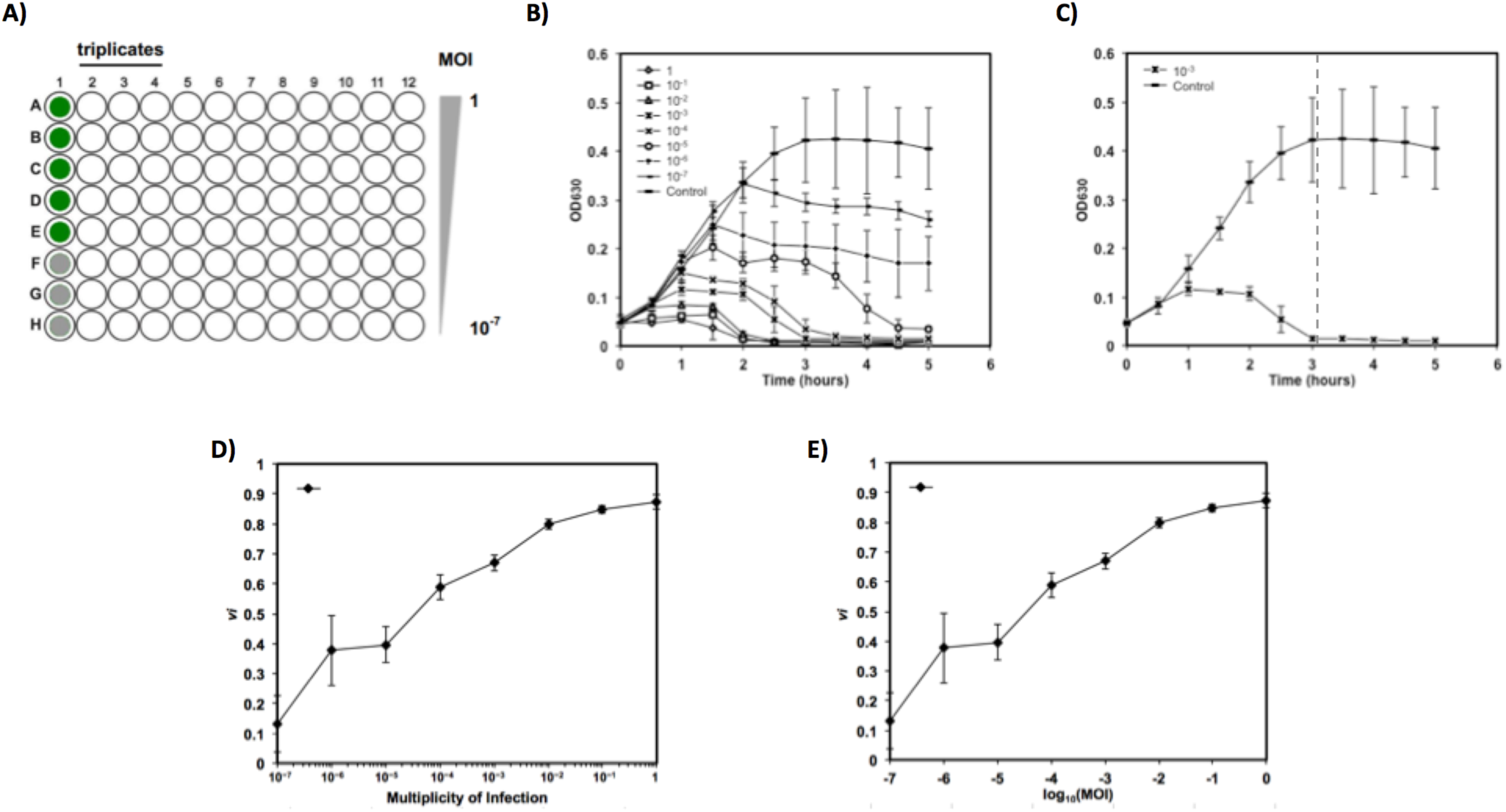
A) Layout of multi-plate for high throughput evaluation of virulence. Column 1 is used for controls: green wells for phage-free cultures, gray wells for media blanks. Each column is used to test a different phage. Rows indicate different initial MOIs, from 1 (row A) to 10^-7^ (row H). **B)** Bacterial reduction curves for phage T4 infecting *E. coli* 11303 in TSB at 37°C. **C)** Growth curve of phage-free control and bacterial reduction curve at MOI 10^-3^ (extracted from B). The dashed vertical line indicates the limit of integration. **D)** Virulence curve generated from bacterial reduction curves of phage T4 growing in TSB at 37°C. Each individual data point represents the local virulence (*v*_*i*_) at the corresponding MOI. Dotted line highlights MV50 determination. **E)** Virulence curve from D) with x-axis converted to log_10_(MOI). Error bars depict the standard deviation of six replicates.

### Evaluation of Phage Cocktails

The virulence of two combinations of phages T5 and T7 (at proportional T5:T7 ratios of 1:1 and 3:1) was also assessed using the same procedure as described above, with the particularity that the MOI reported was the combined MOI of both phages. For example, when the ratio of 1:1 was tested, 0. 5×10^8^ pfu/ml of phage T5 and 0.5×10^8^ pfu/ml of phage T7 were used to make a total combined titer of 1×10^8^ pfu/ml, for an MOI of 1. Similarly, for the 3:1 ratio, 0.75×10^8^ pfu/ml of phage T5 and 0.25×10^8^ pfu/ml of phage T7 were used for a total combined titer of 1×10^8^ pfu/ml. The use of the combined MOI is important for the assessment of virulence in comparison to results with single phages; this allows the rapid identification of synergistic or inhibitory effects between phages. In these tests, the microplate layout described in Fig. 1A was still used, where triplicates of phage cocktails occupied 3 columns. Incubation in TSB at 37°C, measurements and analysis were conducted in the same manner as for single phage testing.

### Statistical Analysis

Virulence assays for each phage species were performed in duplicate on three separate micro-well plates (n=6) or in triplicate in a single plate (n=3) for the comparison of phage cocktails. This was done so that inaccurate readings or potential contamination could be detected. Data points on graphs are shown as the average of all replicates; error bars depict the standard deviation. Errors reported for Virulence Index (*V*_*P*_ – defined below) values are the summation of errors in all the local virulence (*v*_*i*_ – defined below) values from which the *V*_*P*_ was determined.

## Results

### Virulence Protocol and Nomenclature

The virulence measurements presented herein build on the premise of bacterial reduction curves. A set of bacterial reduction curves – performed as described in the section above – for T4 infecting *E. coli* ATCC 11303 in TSB at 37°C is shown in Fig. 1B. The phage-free control exhibits a classic growth pattern. As can be seen in the other curves found in Fig. 1B, the presence of the phage can significantly reduce the presence of bacteria. By comparing the bacterial reduction curves to the control, we can quantify the reduction due to the killing or damaging of the host by the phage. This is done by comparing the integrated area of a bacterial reduction curve (*A*_*i*_, where *i* is the MOI) to the integrated area of the phage-free control (*A*_0_) as shown in Fig. 1C for the MOI 10^-3^. These areas are calculated from the onset of infection (time 0) to the time of the onset of the stationary phase in the phage-free control (indicated by the vertical dashed grey line near 3h in Fig. 1C). It is important to stress how the establishment of the limit of integration plays a significant role in the assessment of virulence. This limit should be set as the onset of stationary phase in the phage-free control (at 3h in Fig. 1C). This provides a consistent reference for integration that can be easily identified for any phage-host system and restricts measurements to the period of cell growth – a necessary condition for productive infection for many phages.^37^ Moreover, it ensures that the range of the virulence measurements is well distributed no virulence and maximum virulence, as discussed below. In general, we recommend establishing the limit of integration at the time when the slope of OD_600_ over time reaches ≤ 0.03 h^-1^.

Using the two areas calculate for the free-phage control (*A*_*o*_) and the culture infected at a given MOI (*A*_*i*_), a local virulence (*v*_*i*_), capturing the dynamics of phage infection, can be calculated for that specific MOI under a given set of conditions:

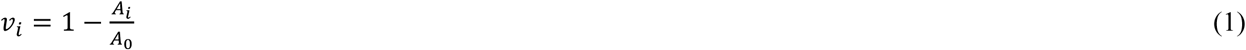

where *v*_*i*_ reports the killing or damaging ability of the phage at a given MOI. The more virulent a phage, the faster it kills a large number of bacteria (or prevents them from growing), and the greater *v*_*i*_. Local virulence is measured on a scale from 0 to 1, where 0 represents the absence of virulence and 1 represents maximum theoretical virulence – instantaneous cell death. While this theoretical maximum is unlikely to be observed in the laboratory, a *v*_*i*_ over 0.90 is readily achievable with the appropriately lytic phage-host system.

Additionally, a virulence curve (Fig. 1E) can be obtained for the phage grown under a given set of environmental conditions (*e.g.* temperature, medium, ionic strength) by plotting local virulences calculated from Fig. 1B against *log MOI* (which is easily obtained from the conversion of the MOI axis seen in Fig. 1D). This provides a powerful tool in characterizing a phage against a specific host over a large range of MOIs. Generally, the earlier *v*_*i*_ approaches 1 on the virulence curve, the more virulent the phage. Two important values can be gathered from such virulence curves to quantify phage virulence: the Virulence Index (*V*_*P*_, where *P* refers to the phage species) and MV_50_ (MOI required to produce a local virulence of 0.5).

The Virulence Index is defined as the area under the virulence curve (*A*_*P*_) divided by the theoretical maximum area under the virulence curve (*A*_*max*_):

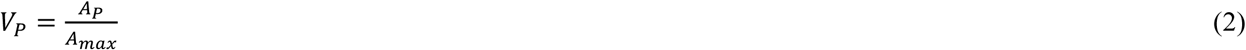

where *A*_*P*_ is determined by integrating the virulence curve according to:

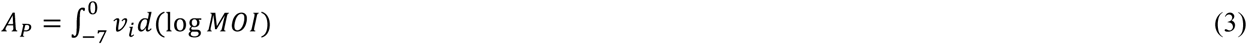

Note that log MOI is used in the integration in order to give equal weighting to each *v*_*i*_ when calculating the Virulence Index. The value for *A*_*max*_ is determined by Eq. 3 under the special condition that *v*_*i*_ = 1 for all MOIs. In this study, *A*_*max*_ = 7 for all conditions tested, since the range of MOIs investigated goes from log MOI = −7 to log MOI = 0.

Similarly to the local virulence, the Virulence Index is normalized such that the theoretical maximum is 1. To reach this value, instantaneous lysis of the entire bacterial culture for all MOIs tested would be required, a physical impossibility. In this study, the highest Virulence Index observed was 0.84. A Virulence Index of 0 signifies the complete absence of virulence over the range of MOIs tested.

From the data shown in Fig. 1D (phage T4 in TSB at 37°C) a Virulence Index of 0.6 was obtained. It was calculated as follows (integrations calculated using the trapezoid rule):

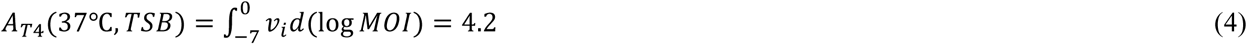

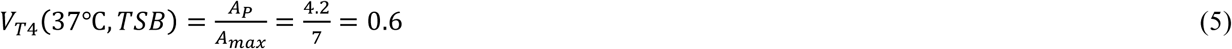

The final quantitative parameter given by the virulence curve is *MV*_*50*_, an analog to *ID*_*50*_ (infective dose for 50% of subjects) used in toxicology.^38^ It is defined as follows:

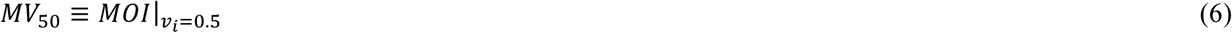

The MV_50_ is the MOI for which the local virulence *v*_*i*_ is equal to 0.5; the MOI at which the phage achieves 50% of the maximum theoretical virulence. This provides another tool for comparing phage virulence. The lower the MV_50_, the more virulent the phage. The MV_50_ can be found through inspection of the virulence curve. As an example, the MV_50_ for phage T4 at 37°C in TSB obtained from Fig. 1D is:

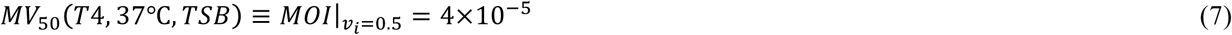

Note that the MV_50_ should be reported to only one significant figure due to the intrinsic error in the virulence curve. This measure is meant as a means of comparison of the order of magnitude of virulence.

### Comparison of Virulence

Bacterial reduction curves and their corresponding virulence curves are shown for phages T5 (Fig. 2A and 2B) and T7 (Fig. 2C and 2D) in TSB at 37°C.

**Figure 2.**
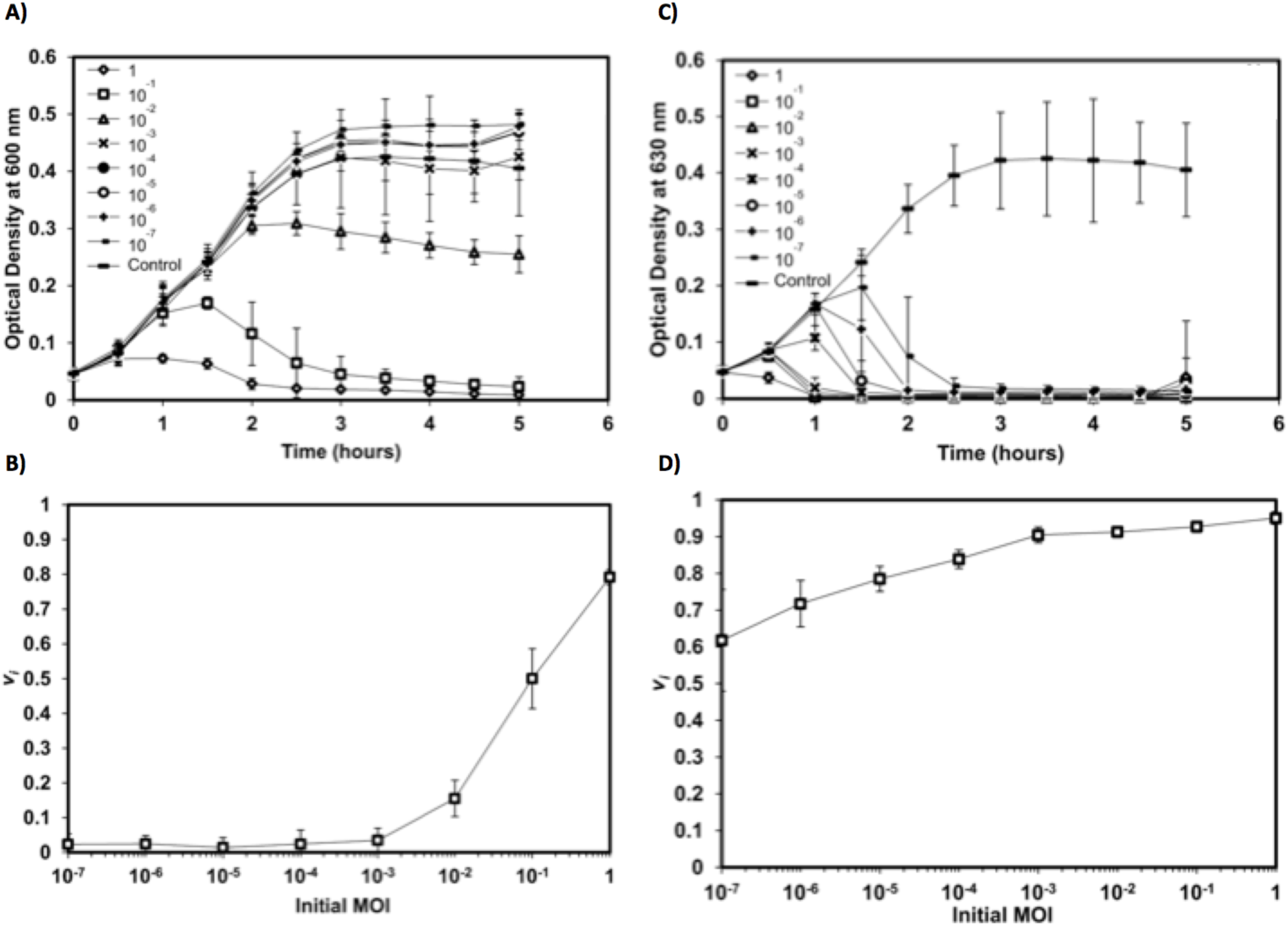
A) Bacterial reduction curves and **B)** corresponding virulence curve for phage T5 infecting *E. coli* 11303 in TSB at 37°C. **C)** Bacterial reduction curves and **D)** corresponding virulence curve for phage T7 infecting *E. coli* 11303 in TSB at 37°C. Error bars depict the standard deviation of six replicates.

Phage T5 was the least virulent phage studied (Fig. 2B). In contrast to the bacterial reduction curves exhibited by phage T4 (Fig. 1D) and phage T7 (Fig. 2D), phage T5 only began to show measurable virulence at MOIs ≥ 10^-2^. For all MOIs below this threshold, the values of local virulence observed were 0 (Fig. 2A). Although being noticeable only from an MOI of 10^-2^, the local virulence increased rapidly, peaking at 0.8 at an MOI of 1. Accordingly, the Virulence Index measured for T5 under the conditions tested was *V*_*T*5_(37°C,*TSB*) = 0.17.

Phage T7 was the most virulent phage tested under these conditions and yielded a Virulence Index of 0.84 (from Fig. 2D). The remarkably virulent nature of this phage is observed qualitatively in the bacterial reduction curves (Fig. 2C). At MOIs ranging from 1 to 10^-3^, culture-wide lysis was achieved within 1 h. Even at the lowest MOI tested (10^-7^) culture-wide lysis was observed after 2.5 h; whereas the phage-free control grew to stationary phase in 3 h (Fig. 2C).

While the virulence curves can be used to quickly and effectively compare different phages, mutants or progeny, they also provide excellent visual tools for analysis and comparison of how a phage behaves under a set of conditions. For example, Fig. 3 compares the virulence curves for phages T4, T5, and T7 in different environmental conditions. It is clear from these curves that temperature (Fig. 3A vs. 3B) and media composition (Fig. 3B vs. 3C) significantly impact phage virulence. Importantly, the same range of MOIs must be used when comparing virulence across different phage species or conditions. If virulence measurements were limited to MOIs ranging from 10^-2^ to 1, inspection of Fig. 3C would lead one to conclude that phages T4 and T7 have very similar behaviours. However, inspection of the entire experimental range demonstrates significant differences in the virulence between these phages.

**Figure 3.**
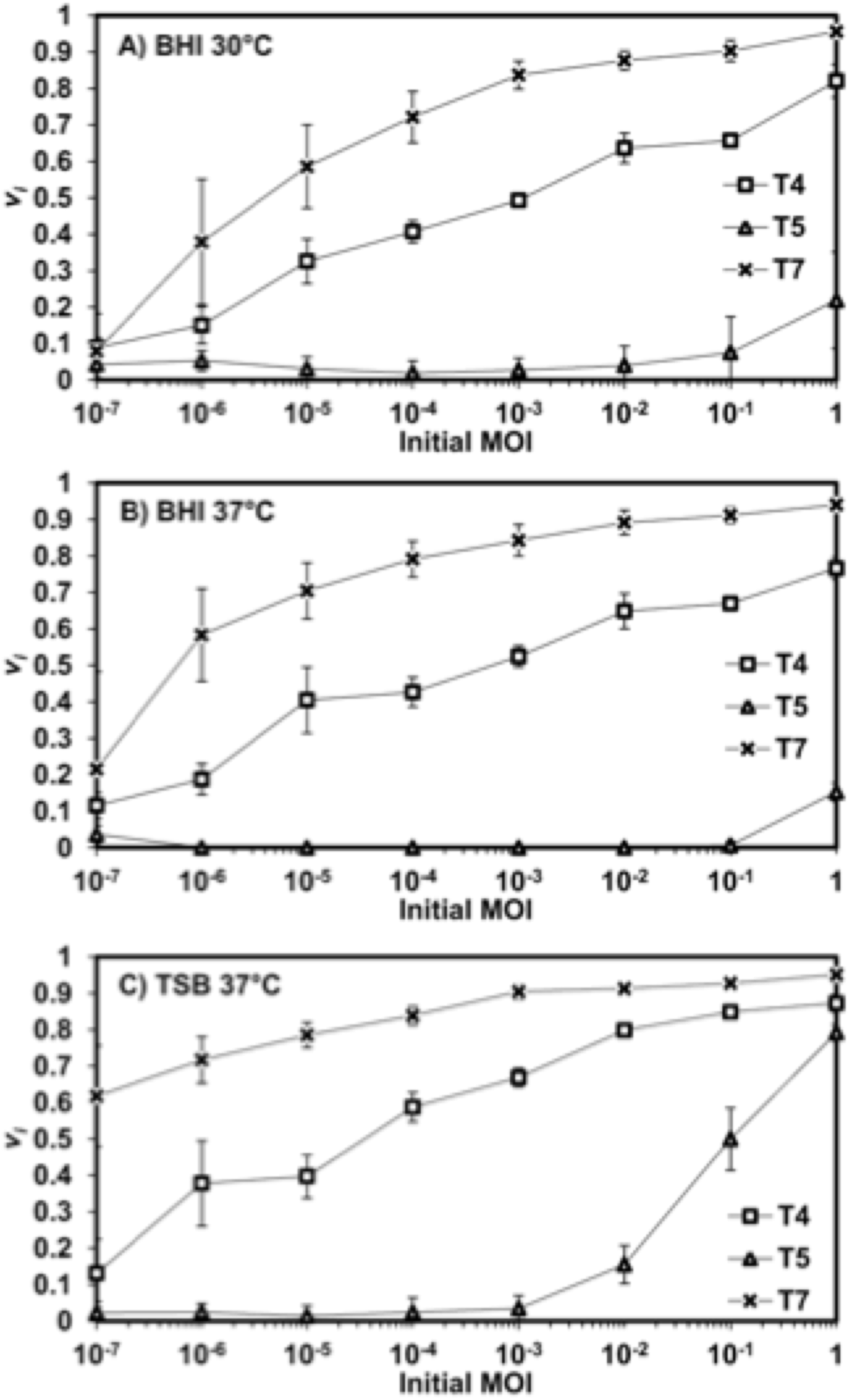
Virulence curves of phages T4, T5, and T7 in **A)** BHI at 30°C, **B)** BHI at 37°C, and **C)** in TSB at 37°C. Error bars depict the standard deviation of six replicates.

Table 1 displays the Virulence Index (*V*_*P*_) and MV_50_ values for phages T4, T5, and T7 in two different media (TSB and BHI) at two different temperatures (30°C and 37°C). The data demonstrates the full spectrum of virulence, from the limited virulence of phage T5 in BHI at 37°C (*V*_*T*5_ = 0.05, *MV*_50_ > 1) to the exceptional virulence of T7 in TSB at 37°C (*V*_*T*7_ = 0.84, *MV*_50_ < 10^-7^). Together the Virulence Index and the MV_50_ values provide a framework for assessing the virulence of a phage and for comparing phage virulence across species and conditions.

**Table 1:** Virulence and *MV*_*50*_ values for phages T4, T5, T7, and combinations of phages T5 and T7 under various conditions.

| Phage | Virulence Index ($V_P$) | | | | $MV_{50}$ | | | |
| --- | --- | --- | --- | --- | --- | --- | --- | --- |
|  | 30° C |  | 37° C |  | 30° C |  | 37° C |  |
|  | TSB | BHI | TSB | BHI | TSB | BHI | TSB | BHI |
| T4 | 0.55 | 0.45 | 0.60 | 0.47 | $1 \times 10^{-4}$ | $1 \times 10^{-3}$ | $4 \times 10^{-5}$ | $6 \times 10^{-4}$ |
| T5 | 0.28 | 0.05 | 0.17 | 0.01± | $5 \times 10^{-1}$ | > 1 | $1 \times 10^{-1}$ | > 1 |
| T7 | 0.77 | 0.69 ± | 0.84 | 0.76 | $8 \times 10^{-7}$ | $6 \times 10^{-7}$ | $< 10^{-7}$ | $4 \times 10^{-6}$ |
| T5:T7<sub>(1:1)</sub> | - | - | 0.81 | - | - | - | $5 \times 10^{-6}$ | - |
| T5:T7<sub>(3:1)</sub> | - | - | 0.72 | - | - | - | $1 \times 10^{-6}$ | - |

Using the values calculated for phages T4, T5, and T7, it is a straightforward task to identify the relative virulence of each phage. Phage T7 had the highest virulence values and lowest MV_50_ values in all conditions tested. Phage T5 consistently had the lowest virulence and highest MV_50_ values. The effect of media conditions on virulence is also apparent. For all phages, BHI unfailingly resulted in lower virulence values. However, the effect of temperature, over the admittedly narrow range tested, was more muted. For phage T4, no significant effect of temperature on virulence was seen between 30°C and 37°C. For T7, virulence increased with temperature, while it decreased for T5.

### Evaluation of combinations of phages

Similarly, the virulence curves for the mixtures of phages T5 and T7 provide information on the performance of combinations of phages or of phage cocktails. As can be observed in Figure 4, the reduction curves of the combinations of phages (ratios of phage T5 to T7 of 1:1 and 3:1) demonstrate a killing potential between those of the single phages, with the 1:1 ratio showing more virulence. This is also observed in the resulting values of virulence index (0.81 and 0.72, respectively) and MV_50_ (5×10^-6^ and 1×10^-6^, respectively). It is interesting to note that the Virulence Indexes obtained are not proportional to the relative initial abundance of each phage in the cocktail. This is consistent with the fact that, in the absence of interference or inhibition between them, the contributions of individual phages in a cocktail depend on their individual infection kinetic parameters.

**Figure 4.**
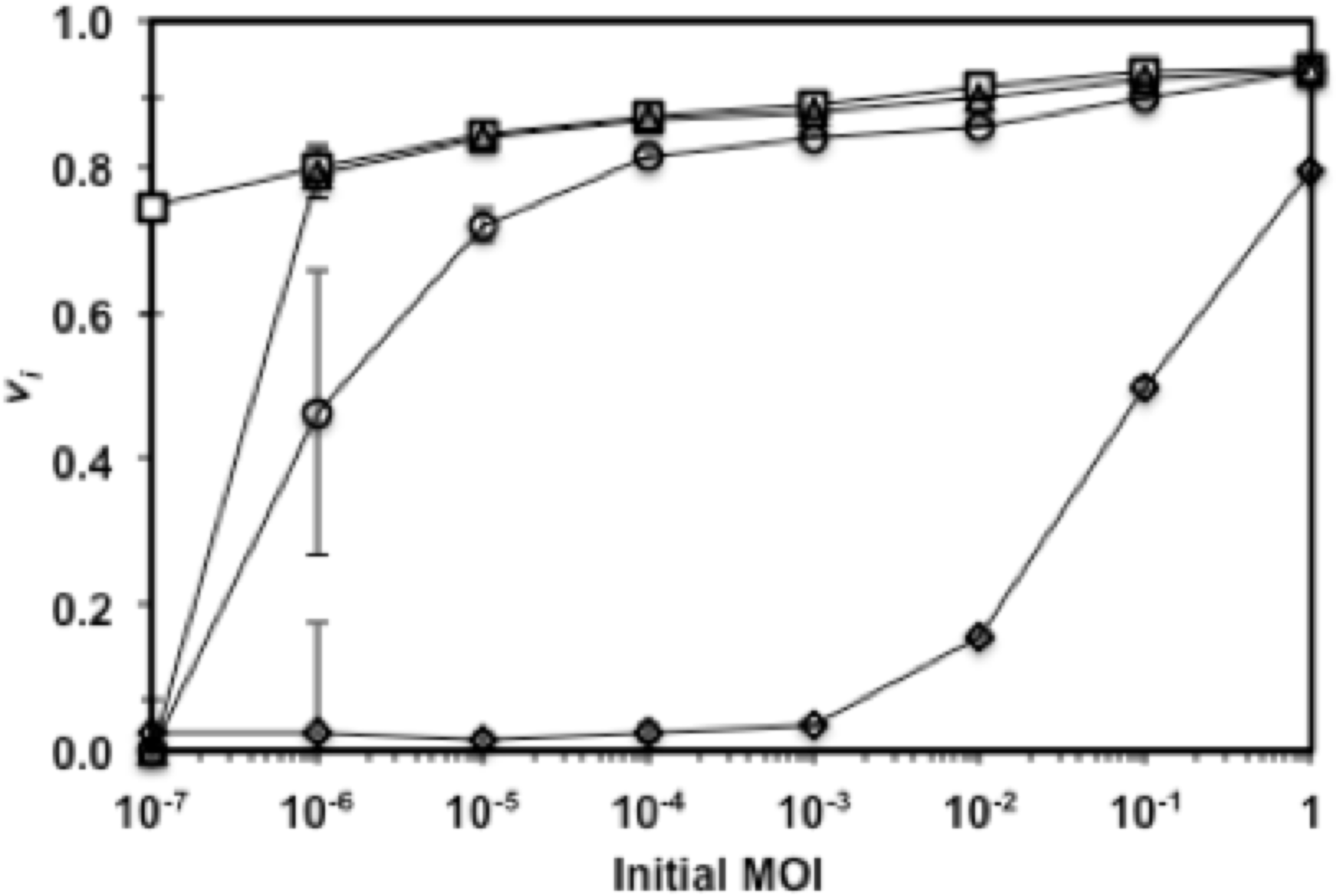
Virulence curves of phages T5 (diamonds), T7 (squares), and combinations of phage T5 and T7 (3:1, circles; 1:1, triangles) in TSB at 37°C. Error bars depict the standard deviation of triplicates.

## Discussion

Unlike most other biologics, phages are intrinsically dynamic in nature, be it through mutations ^39-43^ or phenotypic traits.^43-46^ In addition, infections are influenced by many factors, some correlated, others orthogonal, but all affecting infection dynamics. The evaluation of single parameters (e.g. titer, burst size, efficacy of plating, etc.) is not sufficient. Generally, individual phage characteristics are measured and discussed in relation to how they affect virulence.^19,40,47^ For example, faster adsorption rates, larger burst sizes, and shorter latent periods are all conditions observed to increase phage virulence. But there is no current way of relating these parameters or weighing their individual or combined input towards an overall phage virulence. Hence, the characterization of phages, the evaluation of their efficacy and quality control through the manufacturing process all require a different approach, one that encompasses the effects of all factors influencing virulence.

As mentioned above, researchers have been using qualitative observations of bacterial reduction curves to describe phage virulence for many years. In fact, groups have proposed a cell susceptibility scale based on visual observations of the bacterial reduction curve ^24^ and a primary comparison of the area under reduction curves.^35,36^ The Virulence Index and MV_50_ measurements forego these limitations and enable direct, quantitative comparisons of phage virulence, compounding all the parameters affecting it.

The local virulence and Virulence Index are valuable tools for screening new phage isolates and comparing phages for specific applications. Consider the case of the bacterial reduction curve for phage T7 at an MOI of 10^-7^ (Fig. 2C). It achieved a local virulence value of *v*_10^-7^_ = 0.62, and complete lysis of the culture occurred within 2 hours; astonishingly fast considering such a low MOI. In fact, the virulence curve (Fig. 2D) begins at this value and increases steadily before stabilizing above 0.9 for MOIs ≥ 10^-3^. Therefore, for T7, the MV_50_ is less than 10^-7^. In contrast, for phage T5 under the same conditions, the MV_50_ was 10^-1^ – over 6 orders of magnitude larger than for phage T7. In order to achieve the same lytic capability as one phage T7, more than 1 million phage T5 virions are required at the onset of infection.

An important consideration when comparing local virulence, Virulence Index, and MV_50_ values among various phages and/or conditions is that these parameters are indirect measurements of infection dynamics. Since they are calculated based on the relationship between infected cultures and the growth of the host, they serve as quantitative descriptors of the effect of phages on cell cultures. They are impacted by all the environmental and physiological factors influencing the host and/or phage growth and propagation rates. Thus even if infections may be slower at one set of conditions compared to another (due to slower adsorption rates, longer lysis times, or smaller burst sizes), if the growth of the host is also slowed down by the same magnitude, the virulence may remain the same. This is a powerful aspect of these measurements: they reflect kinetics without being impeded by them.

Phage virulence is thus affected by a large number of environmental and physiological parameters. For example, conditions such as temperature, pH, media composition, and aeration can all affect both host cell growth rate – influencing phage growth-associated parameters such as burst size and latent period – and phage adsorption rate – affecting the rate of infection.^37^ Moreover, phage infections at the population level can exhibit characteristics not easily observed in individual host-phage interactions (e.g. lysis inhibition in the case of T4). All these interacting variables contribute to virulence. A recent study testing the efficacy of six newly isolated phages to treat *Pseudomonas aeruginosa* infection of *Drosophila melanogaster* found no correlation between burst size, adsorption rate, or latent period and phage therapy efficacy.^47^ The only phage parameter found to correlate significantly with treatment efficacy was phage growth rate – implying that a measurement of phage virulence needs to account for contributions of all the disparate factors working together.

With the introduction of the Virulence Index, there now exists a metric which can be used to quantify the pooled contribution of each of these variables on overall phage virulence. Which features of phages T4, T5 and T7 lead to such significant differences in virulence when infecting *E. coli* in TSB? Phage T4 has a very high adsorption rate and adsorption efficiency in TSB ^48^, while T5 and T7 both have similar adsorption rates with poor adsorption efficiencies in the same medium.^49^ Perhaps this poor adsorption efficiency is a major reason why T5 displays limited virulence in TSB. Yet, T7, which has even lower adsorption efficiency, is extremely virulent. In this case, the latent period and/or burst size may be the deciding factors. While not measured in these experiments, T7 is reported to exhibit latent periods of less than 20 minutes while T5’s latent period can exceed 40 minutes.^50^ Note that the relative importance of each of these parameters in determining virulence may also be influenced by the bacterial cell density. The virulence curve can be used in conjunction with traditional measurements of phage growth kinetics to determine the contribution each phage growth parameter has on overall phage virulence.

In addition, as highlighted by many studies ^51-54^, the formulation of phage cocktails is crucial to the success of many phage-based treatments and technologies. Considering most cocktails are composed of more than two phages, proper optimization studies require testing vast numbers of possible formulations, which can be difficult and time-consuming. Hence, it would be most practical to perform a large number of comparisons at a given MOI on a single multi-plate; essentially comparing the values of local virulence (*v*_*I*_) for various combinations of phages. In this case, the whole multi-plate layout could be modified to have various formulations all at the same MOI (rather than the range of MOIs described and shown in Fig. 1). On the other hand, the Virulence Index and the shape of the virulence curve (Fig. 4) can also be used to further understand some of the interactions between the individual phages making up the cocktail. For example, to see if the presence of a phage in the cocktail impedes on the overall virulence.

## Conclusion

The developed methodology will facilitate and standardize the important screening steps used in selecting a phage for specific applications. It can also be used to benchmark different phages or production batches for consistency and quality control purposes, or serve as a reliable comparison platform in the elaboration of formulation of phage cocktails. Finally, it can easily be integrated in high throughput strategies, a non-negligible factor as phage isolation and screening efforts are rapidly expanding.

## Acknowledgements

Funding for this research was provided by the Natural Sciences and Engineering Research Council of Canada, Alberta Innovates Technology Futures and the University of Alberta Research Experience program. Special thanks to Melissa Harrisson who performed the assays for the combinations of T5-T7.

## Authorship Confirmation Statement

ZS and DS designed experiments. ZS, MRT, and KM performed experiments and analyzed data. ZS, MRT, and DS developed virulence metrics. ZS, MRT, and DS wrote the manuscript.

## Author Disclosure Statement

No competing financial interests exist.

